# Chronotype-specific impacts of sleep-wake schedule on behavioral, physiological, and psychological parameters

**DOI:** 10.1101/2025.11.22.684496

**Authors:** Toko Haga, Seitaro Iwama, Junichi Ushiba, Junichi Ushiyama

## Abstract

Diurnal variations in human performance are known to be associated with chronotype, which reflects individual differences in preferred sleep-wake cycles. Early chronotypes (ECs) exhibit superior motor and cognitive functions in the morning, whereas late chronotypes (LCs) tend to perform better in the evening. However, previous studies have examined these patterns under socially imposed nighttime sleep schedules, without considering individuals’ preferred sleep timing. Thus, it remains unclear whether these chronotype-dependent performance peaks persist when individuals follow their preferred sleep schedules. To address this question, we measured subjective states, sleep pressure (using electroencephalography), and motor and cognitive functions at 1.5 and 12.5 hours after awakening under two conditions: the Clock-Time condition (10:30 PM–6:30 AM sleep aligned with social rhythms) and the Self-Time condition (8 hours of sleep aligned with individual preferences). The results revealed distinct chronotype-specific patterns. ECs exhibited increases in fatigue and sleepiness dependent on awake duration, showing typical sleep pressure accumulation. LCs were more affected by sleep timing and showed atypical patterns of sleep pressure accumulation compared with ECs. Motor ability tended to improve in the afternoon in both chronotypes, irrespective of sleep-wake schedule, whereas cognitive performance peaked at different stages of wakefulness - earlier for ECs and later for LCs - independent of the time of day. These findings suggest that chronotype-aligned sleep-wake schedules could help optimize fatigue management, cognitive performance, and motor control. Diurnal variations in performance cannot be attributed to clock time alone but emerge from the combined effects of chronotype, awake duration, and sleep-wake regulation.

**Statement of Significance:** This study provides new insights into the ways in which chronotype and sleep-wake schedules jointly shape behavioral, physiological, and psychological dynamics across the day. Whereas previous research has largely emphasized circadian rhythms, our findings underscore the critical role of actual sleep timing in modulating daily functioning. This integrative perspective extends current understanding of the ways in which individual differences in biological timing and social constraints interact to influence human health and performance. Importantly, the results highlight the need for future studies to disentangle circadian and sleep-wake schedule effects, which could inform personalized strategies for optimizing well-being, productivity, and clinical interventions in disorders related to sleep and circadian misalignment.

## Introduction

Human daily life is characterized by a rhythmic structure that regulates waking, working, and resting in accordance with the diurnal cycle. However, adherence to such socially imposed schedules can lead to a phenomenon known as “social jet lag” [1]. Social jet lag negatively affects various aspects of life, including academic performance [2–6], occupational productivity [7–9], and overall health [10–13], and is considered to result from the misalignment between an individual’s biological rhythms and social rhythms [1]. Thus, social jet lag has emerged as a significant public health and societal concern, given its widespread impact on education, workplace efficiency, and well-being [3,8].

The mismatch between biological and social rhythms is largely attributed to chronotype, which refers to individual differences in sleep-wake cycle preferences. Individuals can be categorized into one of three chronotypes: early chronotype (EC), late chronotype (LC), and intermediate chronotype (IC). These types are regulated by a complex interplay of external factors, such as light exposure [14], and innate factors, such as endogenous oscillator periods [15] and genetic variations in light sensitivity [16–19]. Chronotypes exhibit distinct sleep-wake schedules. ECs generally experience an advancement in their sleep-wake cycle, typically preferring to go to bed early in the evening and wake up early in the morning. In contrast, LCs generally experience a delay in their sleep-wake cycle, typically preferring to stay awake until late at night and waking up later in the morning or early afternoon [20].

In addition to differences in sleep-wake schedules, cognitive and motor functions during wakefulness also show specific within-day variation according to chronotype. Several studies have shown that ECs tend to exhibit optimal cognitive and physical functions earlier in the day, while LCs tend to reach their optimal levels later [21–27]. This chronotype-dependent temporal pattern is also reflected in neurophysiological markers, such as cortical excitability and neural plasticity [23].

However, previous studies of chronotypes have mainly examined the effects of chronotype on diurnal variations in cognitive and motor functions under socially imposed nighttime sleep conditions [21–25]. Consequently, these studies have not fully considered differences in chronotype-based sleep-wake preferences. This raises a key question regarding how motor and cognitive functions, as well as neurophysiological markers, respond when individuals follow their preferred sleep-wake schedules. It is possible that allowing individuals to follow their preferred sleep schedules reduces the effort of waking [28] and promotes optimal functioning from the earliest awake duration, irrespective of chronotype. Alternatively, regardless of preferred sleep timing and the time of day, ECs might always perform better in the earlier stages of awake duration, whereas LCs might perform better in the later stages. To address this question, we investigated within-day variations in subjective states, sleep pressure measured via scalp electroencephalography (EEG), motor learning, and cognitive functions between chronotypes, under two sleep-wake conditions: one aligned with a conventional social schedule and the other aligned with individual chronotype preferences. In both conditions, measurements were conducted with fixed timing relative to each participant’s wake-up time. Thus, only sleep timing differed between experimental conditions (Figure 1). This study revealed how chronotype and sleep-wake schedules jointly shape behavioral, physiological, and psychological dynamics across the day.

**Figure 1:**
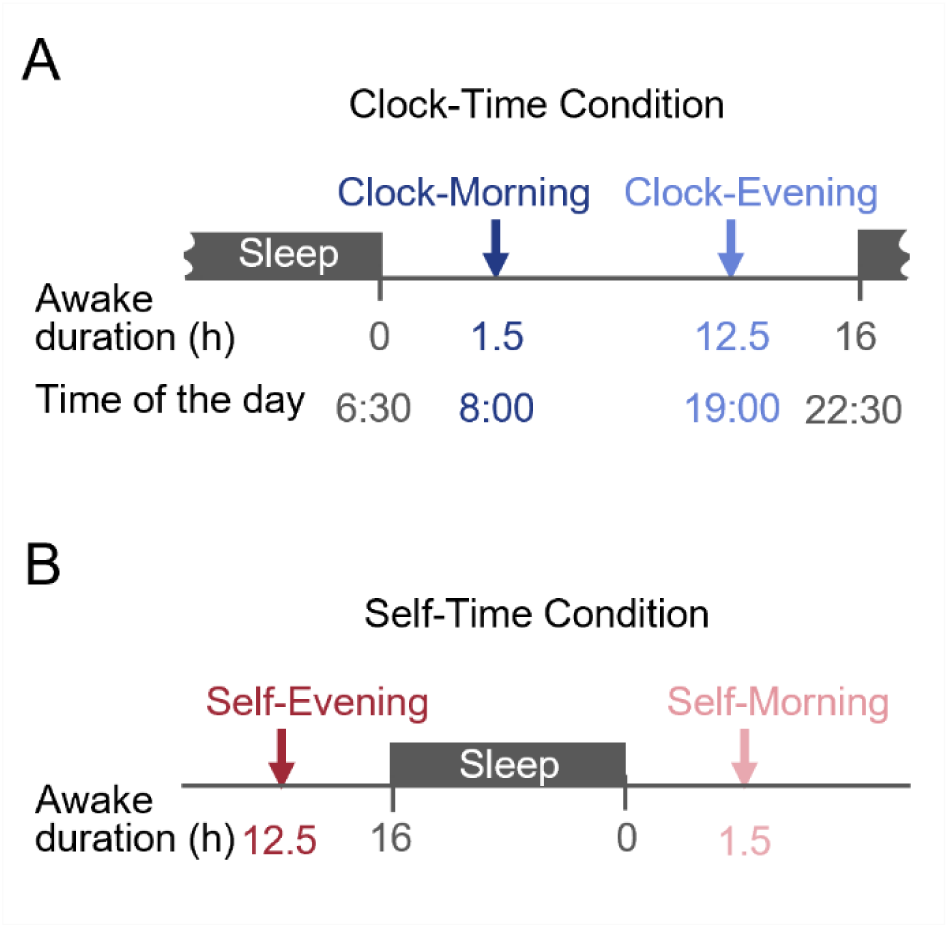
The timing of experimental sessions and sleep schedule under two distinct sleep-wake cycle conditions. **A** The “Clock-Time condition” was aligned with social rhythms. Participants followed a set sleep schedule of 8 hours from 10:30 PM to 06:30 AM (black square). Measurement in the Clock-Morning session was conducted at 08:00 AM,1.5 hours post-awakening. Measurement in the Clock-Evening session was conducted at 07:00 PM, 12.5 hours post-awakening. **B** The “Self-Time condition” was aligned with each participant’s biological rhythm preferences. Participants slept for 8 hours according to their preferred timing (black square). Measurement in the Self-Morning session was conducted 1.5 hours after the participant’s individual waking time. Measurement in the Self-Evening session was conducted 12.5 hours after the participant’s individual waking time.

## Methods

### Participants

We recruited 30 healthy young adults (14 males and 16 females; age: 24.8 ± 2.1 years), who were classified into two chronotype groups: early chronotypes (ECs) and late chronotypes (LCs). Each chronotype group consisted of 15 participants (seven men and eight women). Chronotype was initially assessed in 178 volunteers using the Japanese version of the Morningness-Eveningness Questionnaire (JMEQ) [20]. On the basis of their JMEQ scores, 21 participants were classified as EC, 48 participants as LC, and 109 participants as IC (Figure 2). Subsequently, 15 participants from each group were selected to participate in the main experiment. We balanced participants’ gender and kept the age range to early adulthood (approximately 22–28 years) between the two groups. All participants were right-handed, non-smokers, and passed a screening assessment to confirm the absence of neurological diseases, epilepsy, seizures, medications, metal implants, and current pregnancy. To prevent potential interference with cortical excitability, caffeine and alcohol intake and intense physical activity were prohibited. In addition, sleep duration, daytime activity levels and heart rate were monitored throughout the experimental period using a wearable activity tracker (Fitbit Inspire 2, Fitbit, Inc., USA). Sessions were rescheduled if participants did not meet these eligibility or behavioral control criteria. This study conformed to the Declaration of Helsinki guidelines and was approved by the SFC Research Ethics Committee of Keio University (Approval Number 334). All participants received a detailed explanation of the experiment and provided informed consent prior to participation. Participants were free to discontinue participation in the experiment at any time.

**Figure 2:**
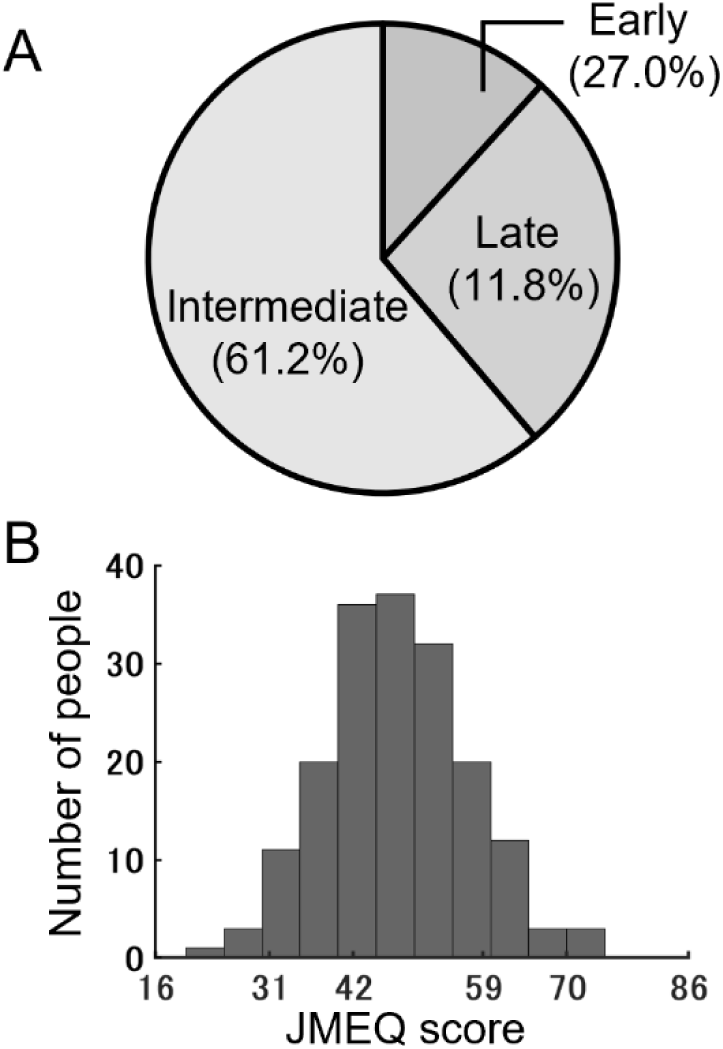
Distribution of chronotypes on the basis of the Japanese version of the Morningness-Eveningness Questionnaire (JMEQ). **A** Pie chart showing the proportion of chronotypes among 178 volunteers: The overall proportions were 27.0% for Early chronotype, 61.2% for Intermediate chronotype, and 11.8% for Late chronotype. **B** Frequency distribution of JMEQ scores. Data are presented on a scale range from 16 to 86. Higher scores indicate stronger morning preference. The score ranges correspond to definite evening type (16–30), moderate evening type (31–41), intermediate type (42–58), moderate morning type (59–69), and definite morning type (70–86) (Ishihara et al., 1986).

### Morningness-Eveningness Questionnaire (MEQ)

The JMEQ [20] was used to identify chronotypes. This questionnaire assesses individuals’ preferred rhythms, indicating their “feeling best” time blocks rather than actual sleep times or the real time for daily/weekly activities (e.g., physical exercise, tests, work). The questionnaire consisted of 19 questions that evaluate morning alertness, morning appetite, and evening tiredness. The questionnaire included 14 items rated on a 4-point scale (score range, 1–4), two items rated on a 5-point scale (score range, 0–4), and three items rated on a 6-point scale (score range, 0–5). Each question is assigned a score, and the total score ranged from 16 to 86. The five chronotype categories were identified by the result s of the JMEQ, as follows: definite evening (16–30), moderate evening (31–41), intermediate or neutral (42–58), moderate morning (59–69), and definite morning (70–86) types. The questionnaire has demonstrated high reliability and a significant correlation with circadian rhythm-related hormonal changes, including melatonin [20].

### Experimental procedure

Before beginning the main experiment session, participants completed a practice session under direct supervision to ensure that they were familiar with the procedure. Participants received instructions about how to wear the EEG device to maintain consistent electrode positioning. Subsequently, they took the EEG device, an activity tracker (Fitbit Inspire 2, Fitbit, Inc., USA), and the personal computer (PC) to their homes for a loan period of approximately 1 week. During this period, participants completed four experimental sessions, structured as a 2 (conditions: Clock-Time vs. Self-Time) × 2 (sessions: Morning vs. Evening) design, with the order of conditions randomly assigned.

In the Clock-Time condition, participants followed a socially aligned sleep schedule of 8 hours, from 10:30 PM to 06:30 AM. Measurements were taken at two fixed times: 08:00 AM (Clock-Morning session, 1.5 hours post-awakening) and 07:00 PM (Clock-Evening session, 12.5 hours post-awakening) [23,26,29] (Figure 1A). The timing of sleep and measurements at the Clock-Time condition was determined on the basis of the average of sleep times and level of activity of ECs and LCs reported in previous studies [26,29]. For the Clock-Morning session, the start time (08:00 AM) was directly adopted from prior chronotype studies [23], which used this timing to represent typical school and work start times. The Clock-Evening session (07:00 PM) was specified on the basis of the time at which both chronotypes were reported to exhibit a comparable level of activity in previous studies [23,29].

In the Self-Time condition, participants followed an individualized sleep schedule of 8 hours according to their preferred timing. Measurements were taken at two relative times, 1.5 hours post-awakening (Self-Morning session) and 12.5 hours post-awakening (Self-Evening session), based on each participant’s wake-up time. The timing of sleep and measurements in the Self-Time condition were determined on the basis of the length of sleep and the duration of wakefulness from awakening to each session in the Clock-Time condition (Figure 1B).

In each session, subjective fatigue and sleepiness were measured using questionnaires, brain activity during the resting state was measured using EEG, and motor and cognitive function were measured using behavioral assessments. Throughout the experiment, sleep-wake timing, sleep duration, physical activity, and weather conditions were controlled or accounted for.

### Subjective fatigue and sleepiness

#### Visual analogue scale to evaluate fatigue severity (VAS-F)

Participants rated their physical fatigue and mental fatigue using two visual analogue scales to evaluate fatigue severity (VAS-F) displayed horizontally on a computer screen. Each scale ranged from 0 (no fatigue) to 100 (extreme fatigue), with the slider bar initially positioned at the midpoint (50). Participants evaluated their own current state by adjusting the slide bar using a mouse [30] (Figure 3A).

**Figure 3:**
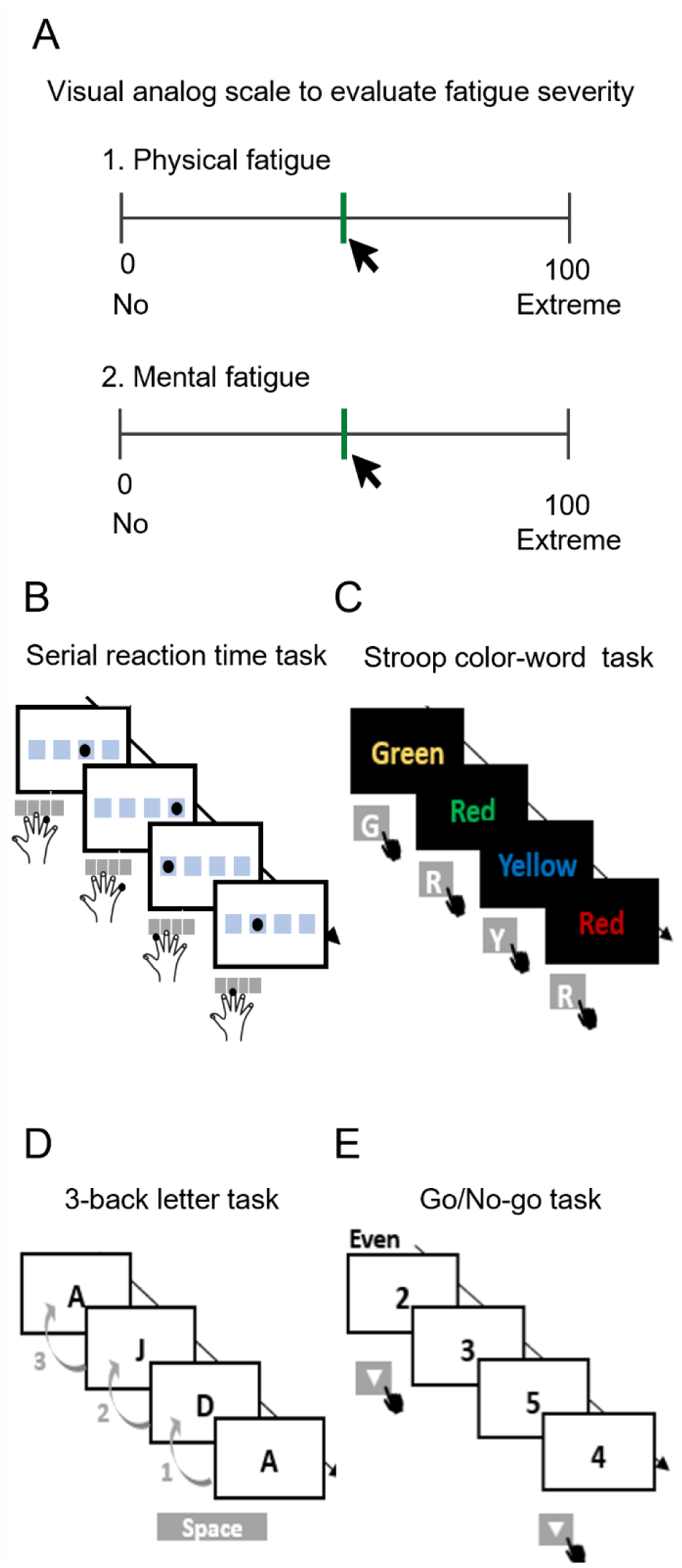
The experiment for each session. **A** Visual analogue scale was used to evaluate fatigue severity (VAS-F). In each session, participants first completed the VAS-F. The scale was presented horizontally on a computer screen with two separate domains: 1) Physical fatigue and 2) Mental fatigue. Each scale ranged from 0 (no fatigue) to 100 (extreme fatigue). The slider bar was initially positioned at 50, the midpoint of the scale, and participants adjusted it using a mouse to reflect their current state. **B** The serial reaction time task (SRTT) was used as a motor learning task. A visual stimulus (a black dot) was presented at one of four horizontally arranged positions on a computer screen, with only one position illuminated at a time. Participants pressed the button corresponding to the dot’s position using the corresponding finger of their right hand as quickly and accurately as possible. **C** The Stroop color-word task was used as a cognitive task. The visual stimuli comprised words displayed in one of four font colors. The task included two types of trials: “congruent” trials, in which the font color matched the word meaning, and “incongruent” trials, in which the font color differed from the word meaning. Participants pressed the corresponding button on the personal computer (PC) keyboard according to the meaning of the word presented (G, R, Y, or B for “green,” “red,” “yellow,” and “blue,” respectively). **D** The 3-back letter task was used as a cognitive task. Visual stimuli consisting of black letters were presented on a computer screen. Participants pressed a shift key on the PC keyboard when the letter on the screen matched the letter presented three trials prior. **E** The Go/No-go task was used as a cognitive task. In two sessions, odd numbers were designated as “Go” stimuli and even numbers were designated as “No-go” stimuli. In the other two sessions, this assignment was reversed, with even numbers serving as “Go” stimuli and odd numbers serving as “No-go” stimuli. Participants pressed a ↓ cursor key on the PC keyboard with the right index finger as quickly and accurately as possible when a “Go” stimulus appeared and withheld their response for a “No-go” stimulus.

#### Karolinska Sleepiness Scale (KSS)

Subjective sleepiness of participants and their alertness were evaluated with the Japanese version of the Karolinska Sleepiness Scale (KSS) [31], which measures sleepiness using a 1–9 Likert-type scale. On this scale, a score of 1 corresponds to “extremely alert” and a score of 9 corresponds to “very sleepy.” Participants were instructed to select the number that best reflected their current level of sleepiness, on the basis of the corresponding descriptions presented on the screen. Participants reported the numerical value that best represented their current state, referring to the values and description of sleepiness presented on the screen.

### Electroencephalography (EEG)

EEG data were continuously recorded during both resting-state and task performance using a telemetry EEG system (AE120E, Nihon Kohden) in combination with a custom -made headphone-type EEG electrode holder. EEG signals were acquired from three dry electrodes positioned at Cz, C3, and C4, with the reference electrode positioned at the mastoid and the ground electrode located on the right cheek according to the international 10 –20 system with a sampling rate of 200 Hz. Each electrode in the system was arranged with contacts positioned both centrally and peripherally. This method was designed to be user-friendly, enabling participants to easily place the electrodes on the scalp by themselves. To ensure optimal signal quality, participants monitored real-time skin-electrode impedance values displayed on a computer screen prior to the start of the recording and adjusted the electrodes to maintain impedances below 50 k Ω. EEG data acquisition was conducted in a quiet home environment.

### Behavioral measurements: motor and cognitive tasks

Participants performed a behavioral task to measure motor learning, which critically involves the primary motor cortex, including long-term potentiation-like plasticity in this region [23] (Figure 3B). In addition, participants performed three cognitive tasks to monitor working memory, attentional functioning, and cognitive functioning (Figure 3C–E). All tasks were designed using MATLAB software (The MathWorks, Inc., USA) following the protocols and evaluation index established in previous research [23,32].

#### Motor learning task: Serial reaction time task (SRTT)

The serial reaction time task (SRTT) was used for measuring motor sequence learning. Performance on this task is considered to be associated with increased neural activity and cortical excitability of the motor, premotor, and supplementary motor areas, and early learning primarily affects the primary motor cortex [33,34,35]. A visual stimulus (a black dot) was presented at one of four horizontally arranged box positions (A, B, C, or D) on a computer screen. The response buttons were constructed by labeling the 7, 8, 9, and 0 keys on the PC keyboard A, B, C, and D, respectively, to correspond to the box positions. Participants were instructed to press the button corresponding to the dot’s position with the respective finger of their right hand (index finger for A, middle finger for B, ring finger for C, and little finger for D) as quickly and accurately as possible. Participants did not know the order in which the buttons would be presented. The task consisted of seven blocks of 100 trials each (700 trials in total). In blocks 1 and 6, the sequence of dots followed a pseudorandom order such that dots were presented equally frequently in each position and never in the same position in two subsequent trials. In blocks 2–5 and 7, the sequence of dots followed a fixed 10 -item order, which was repeated 10 times (e.g., A-B-A-D-B-C-D-A-C-B). At the end of each stimulus-response pair, there was a 500 ms delay before the next cue was presented. Reaction times (RTs) from stimulus onset to button press were recorded for all trials (Figure 3B).

#### Cognitive tasks: Stroop color-word task

The Stroop color-word task is a neuropsychological test that has been extensively used for measuring selective attention, cognitive inhibition, and information processing speed [36,37]. We conducted a computerized Stroop color-word test similar to the Victoria version, on the basis of previous studies [23,37]. The visual stimuli comprised words (red, green, yellow, or blue) displayed in one of these four font colors. The task included two types of trials: “congruent” trials, in which the font color matched the word meaning (e.g., the word “red” displayed in red font), and “incongruent” trials, in which the font color differed from the word meaning (e.g., the word “red” displayed in blue font). Each session consisted of 120 trials, including 30 congruent and 90 incongruent trials. Stimuli were presented on a screen with a black background for 2,000 ms with an interstimulus interval of 500 ms. Participants were instructed to press the corresponding button on the PC keyboard with their right index finger as quickly and accurately as possible, according to the meaning of the word presented. For example, if the word “red” appeared, participants were required to press the button labeled “R” to indicate the word’s meaning, regardless of the font color (red, r: green, g: yellow, y: blue, b). RTs from stimulus onset to button press were recorded for all trials and analyzed separately for congruent and incongruent trials (see “Data Analysis”) [23] (Figure 3C).

#### Cognitive tasks: 3-back letter task

The 3-back letter task was conducted to measure working memory. In this task, participants indicated whether a letter presented on the screen (the “target letter”) matched the letter presented “*n*” trials earlier (the “cue letter”) [38]. The present study employed a 3-back version of the task [23,39], meaning that participants compared each current letter with the letter presented three positions earlier in the sequence. “Hits” (correct responses) were defined as correct identifications of matching letters. Each session consisted of 123 trials. Visual stimuli consisted of black letters (random sequences of 10 letters [A–J]) presented on a computer screen with a white background for 200 ms with an interstimulus interval of 1,800 ms. Participants were instructed to press a shift key on the PC keyboard with their right index finger as quickly and accurately as possible when a match was detected. Accuracy was measured and RTs from stimulus onset to button press were recorded only whe n the participant pressed the button [23] (Figure 3D).

#### Cognitive tasks: Go/No-go task

The Go/No-go task was conducted for measuring attentional functioning including sustained and transient attention, as well as cognitive and impulse inhibition [32]. Visual stimuli consisted of black numbers randomly selected from 0 to 9 in equal proportions and presented on a white background. The assignment of “Go” and “No-go” stimuli varied across four sessions. In two sessions, odd numbers were designated as “Go” stimuli and even numbers were designated as “No-go” stimuli. In the other two sessions, this assignment was reversed, with even numbers serving as “Go” stimuli and odd numbers as “No-go” stimuli. The order of these assignments was randomized for each participant. Each session included 160 trials, comprising 80 “Go” and 80 “No-go” trials. Stimuli were presented for 40 ms with 1,000 ms interstimulus interval. Participants were instructed to press a ↓ cursor key on the PC keyboard with the right index finger as quickly and accurately as possible when a “Go” stimulus appeared and to withhold responses for a “No-go” stimulus. Accuracy was measured and RTs from stimulus onset to button press were recorded only when the participants pressed the button [32,40] (Figure 3E).

### Data analyses

The PC used for all measurements and data processing was equipped with an 11th Gen Intel(R) Core(TM) i7-11370H @ 3.30 GHz CPU and 16 GB of RAM. Response times were recorded with a precision of 0.0046 ms to ensure accurate latency detection.

#### Subjective fatigue and sleepiness

In the VAS-F, participants’ self-reported scores were determined as the distance in units from 0 to the slider’s final position, representing the participant’s subjective rating of physical and mental fatigue.

In the KSS, participants’ self-reported scores were used directly as numerical indicators of their perceived level of sleepiness.

#### EEG

Raw EEG data recorded from the Cz, C3, and C4 electrodes were preprocessed using a custom MATLAB script (The MathWorks, Inc., USA). To correct for baseline drift, the raw signals were linearly detrended, followed by the application of a notch filter (49 –51 Hz) to remove power line interference. Athird-order Butterworth band-pass filter (1–99 Hz) was then applied to attenuate low- and high-frequency noise. For spectral analysis, the preprocessed signals were analyzed using Welch’s method with 1-second Hanning windows and no overlap, providing an averaged estimate of the power spectral density.

Relative θ-band power was calculated as the ratio of sum of power spectral density values within 4–7 Hz to that within 4–50 Hz. In accord with previous studies, the θ-band power during the eyes-open condition was adopted as an indicator of sleep pressure, reflecting the homeostatic sleep drive [23,41–44]. We focused exclusively on θ-band activity in the current study, which is widely recognized as a robust marker of sleep pressure.

#### Behavioral measurements: motor and cognitive tasks

In the SRTT task, RTs were analyzed for all blocks. The mean and standard deviation of RTs were calculated for each block individually. Subsequently, the difference between the mean RT of block 5 (sequence order) and block 6 (random order) was computed, representing motor learning acquisition. This measure reflects the participant’s response to sequence learning compared with sequence-independent performance. Additionally, the difference between the mean RT of block 6 (random order) and block 7 (sequence or der) was calculated to assess learning retention, indicating the extent to which the participant retained the learned sequence [23].

In the Stroop color-word task, RTs were analyzed for correct responses in all trials, congruent trials, and incongruent trials, separately. The RTs in congruent trials were analyzed when the font color matched the word meaning. The RTs in incongruent trials were analyzed when the font color differed from the word meaning. The mean RTs for correct responses were calculated [23].

In the 3-back letter task, the hit rate and sensitivity index d (d prime) were calculated. The hit rate represents the proportion of correct responses (hits) relative to the total number of trials, while d accounts for both the hit rate and the false alarm rate, which represents the proportion of hits minus the proportion of false-alarms. The mean RTs for correct responses (hits) were calculated [23].

In the Go/No-go task, the overall hit rate and the inhibition rate were calculated. The overall hit rate represents the proportion of correct responses (hits) relative to the total number of trials. The hit accuracy for “Go” trials was calculated as the proportion of correct responses relative to the total number of “Go” trials. Additionally, the inhibition rate was calculated as the proportion of correct responses for “No-go” trials relative to the total number of “No-go” trials. The mean RTs for correct responses (hits) were calculated [32].

#### Statistics

Statistical analyses were performed using SPSS Statistics version 29 (IBM SPSS Inc., Armonk, New York, USA). Normality of the data distribution was assessed prior to the main analyses. For variables that did not meet the assumption of normality, the aligned rank transform procedure was applied before conducting a two-way repeated-measures analysis of variance (rmANOVA) [45]. For variables with normally distributed data, a two-way rmANOVA was directly conducted without transformation. A two-way rmANOVA was conducted to examine the main effects of condition (Clock-Time condition and Self-Time condition) and experiment timing (Morning [1.5 hours after waking up] and Evening [12.5 hours after waking up]) on each evaluation index for ECs and LCs. Additionally, the interaction between condition and experiment timing was analyzed using a two-way rmANOVA, and the F values for the interaction were obtained. When a significant interaction effect was detected, simple main effects were examined, followed by Bonferroni-corrected post hoc tests to adjust for multiple comparisons. For all statistical analyses, a significance level of P < 0.05 was adopted. For post hoc comparisons, Bonferroni correction was applied to control for multiple testing, and only P values below the corrected threshold were considered to be statistically significant.

## Results

### Subjective fatigue and sleepiness

#### VAS-F

To examine whether subjective fatigue varied with the time of day and sleep-wake alignment, subjective physical and mental fatigue were assessed using the VAS-F, with lower scores indicating less fatigue. For ECs, regarding physical fatigue, the two-way rmANOVA results revealed a significant effect of experiment timing (*F* = 5.971, *p* = 0.030, 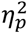 = 0.315, Figure 4, Table 3). The scores in the Morning were significantly lower compared with those in the Evening (Figure 4, Table 1). In contrast, regarding mental fatigue, the two-way rmANOVA results revealed no significant main effects or interaction (Table 3). This finding indicated that there was no significant difference between Morning and Evening or between Clock- and Self-Time conditions in ECs (Figure 4, Table 1). For LCs, regarding physical fatigue, the two-way rmANOVA results revealed a significant effect of condition (*F* = 5.133, *p* = 0.040, 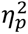 = 0.268, Figure 4, Table 4). The scores under the Self-Time condition were significantly lower compared with those under the Clock-Time condition (Figure 4, Table 2). In contrast, regarding mental fatigue, the two-way rmANOVA results revealed no significant main effects or interaction (Figure 4, Table 4). This indicates that there was no significant difference between Morning and Evening or between the Clock- and Self-Time conditions in LCs (Figure 4, Table 2). Collectively, physical fatigue was associated with awake duration in ECs, whereas in LCs it was associated with the sleep-wake schedule. Mental fatigue showed no significant differences across experiment timing or conditions.

**Figure 4:**
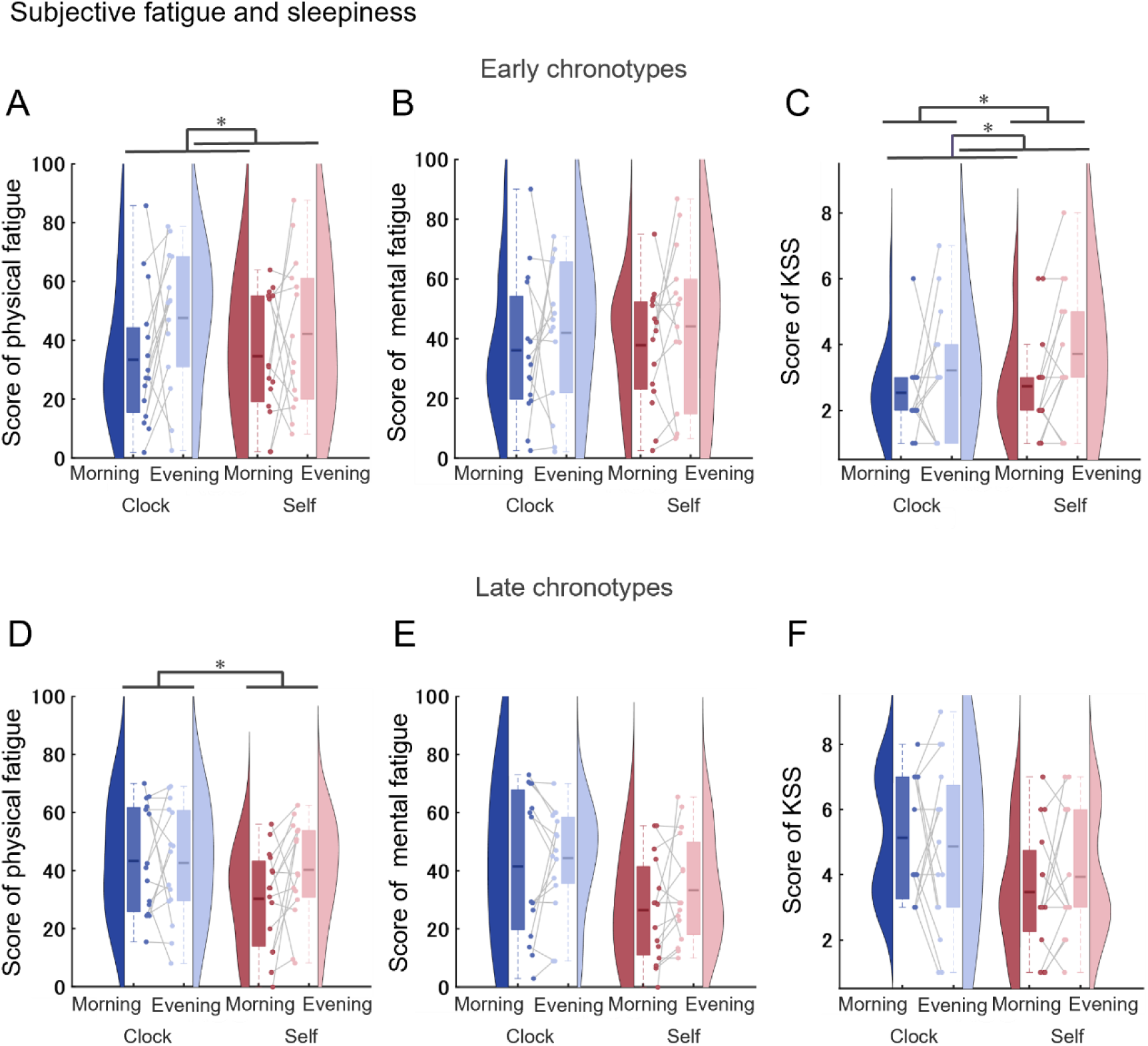
Evaluation of fatigue and sleepiness using visual analogue scale (VAS-F) and Karolinska Sleepiness Scale (KSS) in early and late chronotypes. Physical and mental fatigue severity and sleepiness were measured using the VAS-F and KSS under Clock-Time conditions (Clock-Morning and Clock-Evening) and Self-Time conditions (Self-Morning and Self-Evening) in early chronotypes (ECs) and late chronotypes (LCs). The top panels (A–C) represent data for ECs, whereas the bottom panels (D–F) represent data for LCs. For all measures, lower scores indicate less fatigue or sleepiness**. A, D** Physical fatigue scores from the VAS-F. **B, E** Mental fatigue scores from the VAS-F. **C, F** Sleepiness scores from the KSS. **A–F** were analyzed using a two-way repeated-measures analysis of variance to examine the effects of condition and experiment timing and interaction between condition and experiment timing on fatigue severity (VAS-F) and sleepiness (KSS). Significant main effects of condition and experiment timing are indicated by asterisks (* for *p* < 0.05). Violin plots show the distribution of data. Boxplots show the mean and interquartile range. Dots and connecting lines indicate individual participant data.

**Table 1:** Results of mean± standard deviation for early chronotypes.

| Measurement | Task | Evaluation index | Clock-Time condition |  | Self-Time condition |  |
| --- | --- | --- | --- | --- | --- | --- |
|  |  |  | Morning | Evening | Morning | Evening |
| Subjective fatigue | VAS-F | Score of physical fatigue | 31.40 ± 22.67 | 47.62 ± 23.68 | 34.87 ± 21.68 | 42.23 ± 25.77 |
| | | Score of mental fatigue | 34.39 ± 23.89 | 41.95 ± 24.16 | 38.30 ± 20.60 | 44.16 ± 26.96 |
| Sleepiness | KSS | Score | 2.50 ± 1.22 | 3.21 ± 1.89 | 2.71 ± 1.64 | 3.71 ± 2.09 |
| EEG | EEG | Theta power ratio at Cz | 32.48 ± 8.34 | 39.93 ± 11.68 | 35.78 ± 8.88 | 40.76 ± 12.31 |
| | | Theta power ratio at C3 | 30.85 ± 12.14 | 42.65 ± 14.03 | 34.31 ± 10.15 | 40.75 ± 13.78 |
| | | Theta power ratio at C4 | 29.38 ± 8.40 | 38.79 ± 12.30 | 33.12 ± 11.11 | 40.52 ± 12.65 |
| Motor learning | SRT | Amount of acquisition | 69.36 ± 79.31 | 108.53 ± 97.25 | 79.02 ± 85.79 | 100.73 ± 106.72 |
| | | Amount of retention | 26.90 ± 81.26 | 52.97 ± 74.74 | 53.70 ± 68.73 | 38.42 ± 67.79 |
| Cognitive function | Stroop | Mean RT at overall | 682.62 ± 95.13 | 676.76 ± 103.75 | 696.60 ± 121.47 | 683.30 ± 127.87 |
| | | Mean RT at congruent | 669.86 ± 94.82 | 666.21 ± 104.77 | 688.26 ± 112.63 | 675.85 ± 144.87 |
| | | Mean RT at incongruent | 686.88 ± 96.01 | 680.30 ± 104.11 | 699.43 ± 125.05 | 685.79 ± 123.41 |
| | 3-back | Accuracy at hit | 70.48 ± 18.11 | 70.45 ± 24.19 | 61.43 ± 20.54 | 67.38 ± 25.26 |
| | | Accuracy at d prime | 66.90 ± 17.71 | 67.98 ± 25.15 | 58.33 ± 21.07 | 64.44 ± 25.86 |
| | | Mean RT | 730.52 ± 154.08 | 712.96 ± 142.43 | 675.62 ± 128.81 | 678.35 ± 165.53 |
| | Go/No-go | Accuracy at overall | 93.71 ± 2.94 | 93.21 ± 3.87 | 94.91 ± 2.89 | 91.12 ± 4.70 |
| | | Accuracy at hit | 93.13 ± 5.45 | 92.68 ± 9.13 | 94.91 ± 5.73 | 88.39 ± 8.54 |
| | | Accuracy at inhibition | 94.29 ± 4.85 | 93.75 ± 4.52 | 94.91 ± 4.74 | 93.84 ± 4.29 |
| | | Mean RT | 479.07 ± 47.57 | 462.11 ± 56.94 | 479.73 ± 61.46 | 482.36 ± 68.06 |

*Note.* VAS-F = visual analogue scale to evaluate fatigue severity; KSS = Karolinska sleepiness scale; EEG = electroencephalogram; SRT = serial reaction time; RT = reaction time.

**Table 2:** Results of mean± standard deviation for late chronotypes.

| Measurement | Task | Evaluation index | Clock-Time condition |  | Self-Time condition |  |
| --- | --- | --- | --- | --- | --- | --- |
|  |  |  | Morning | Evening | Morning | Evening |
| Subjective fatigue | VAS-F | Score of physical fatigue | 43.35 ± 18.68 | 42.64 ± 19.81 | 30.34 ± 17.18 | 40.30 ± 16.97 |
| | | Score of mental fatigue | 41.51 ± 25.26 | 44.40 ± 17.98 | 26.49 ± 18.01 | 33.35 ± 18.31 |
| Sleepiness | KSS | Score | 5.13 ± 1.89 | 4.87 ± 2.56 | 3.47 ± 1.88 | 3.93 ± 1.94 |
| EEG | EEG | Theta power ratio at Cz | 36.23 ± 10.50 | 35.57 ± 13.37 | 33.99 ± 13.17 | 32.65 ± 14.97 |
| | | Theta power ratio at C3 | 34.66 ± 11.47 | 35.29 ± 11.35 | 34.99 ± 16.51 | 30.72 ± 13.37 |
| | | Theta power ratio at C4 | 33.43 ± 10.26 | 32.99 ± 13.65 | 32.26 ± 12.90 | 30.13 ± 12.54 |
| Motor learning | SRT | Amount of acquisition | 80.26 ± 96.16 | 128.29 ± 111.84 | 147.87 ± 113.36 | 115.49 ± 108.55 |
| | | Amount of retention | 61.78 ± 97.40 | 83.61 ± 101.83 | 95.69 ± 87.49 | 86.98 ± 88.93 |
| Cognitive function | Stroop | Mean RT at overall | 670.99 ± 121.65 | 635.13 ± 114.98 | 669.36 ± 117.68 | 630.13 ± 102.66 |
| | | Mean RT at congruent | 655.39 ± 123.89 | 625.62 ± 120.62 | 658.35 ± 122.31 | 619.90 ± 117.64 |
| | | Mean RT at incongruent | 670.99 ± 121.65 | 635.13 ± 114.98 | 669.36 ± 117.68 | 630.13 ± 102.66 |
| | 3-back | Accuracy at hit | 59.11 ± 15.40 | 66.67 ± 19.47 | 62.67 ± 16.77 | 60.67 ± 20.52 |
| | | Accuracy at d prime | 54.30 ± 16.07 | 62.15 ± 21.03 | 55.78 ± 17.55 | 60.30 ± 20.71 |
| | | Mean RT | 858.82 ± 164.47 | 830.28 ± 221.39 | 823.77 ± 170.39 | 849.36 ± 220.74 |
| | Go/No-go | Accuracy at overall | 94.67 ± 4.40 | 94.00 ± 4.10 | 95.50 ± 4.27 | 94.25 ± 4.53 |
| | | Accuracy at hit | 95.17 ± 6.28 | 90.67 ± 7.67 | 94.67 ± 6.28 | 95.25 ± 5.27 |
| | | Accuracy at inhibition | 94.167 ± 1.31 | 97.333 ± 0.68 | 96.33 ± 5.76 | 93.25 ± 7.38 |
| | | Mean RT | 465.56 ± 54.81 | 469.95 ± 42.77 | 454.53 ± 46.35 | 469.22 ± 46.14 |

*Note.* VAS-F = visual analogue scale to evaluate fatigue severity; KSS = Karolinska sleepiness scale; EEG = electroencephalogram; SRT = serial reaction time; RT = reaction time.

**Table 3:**
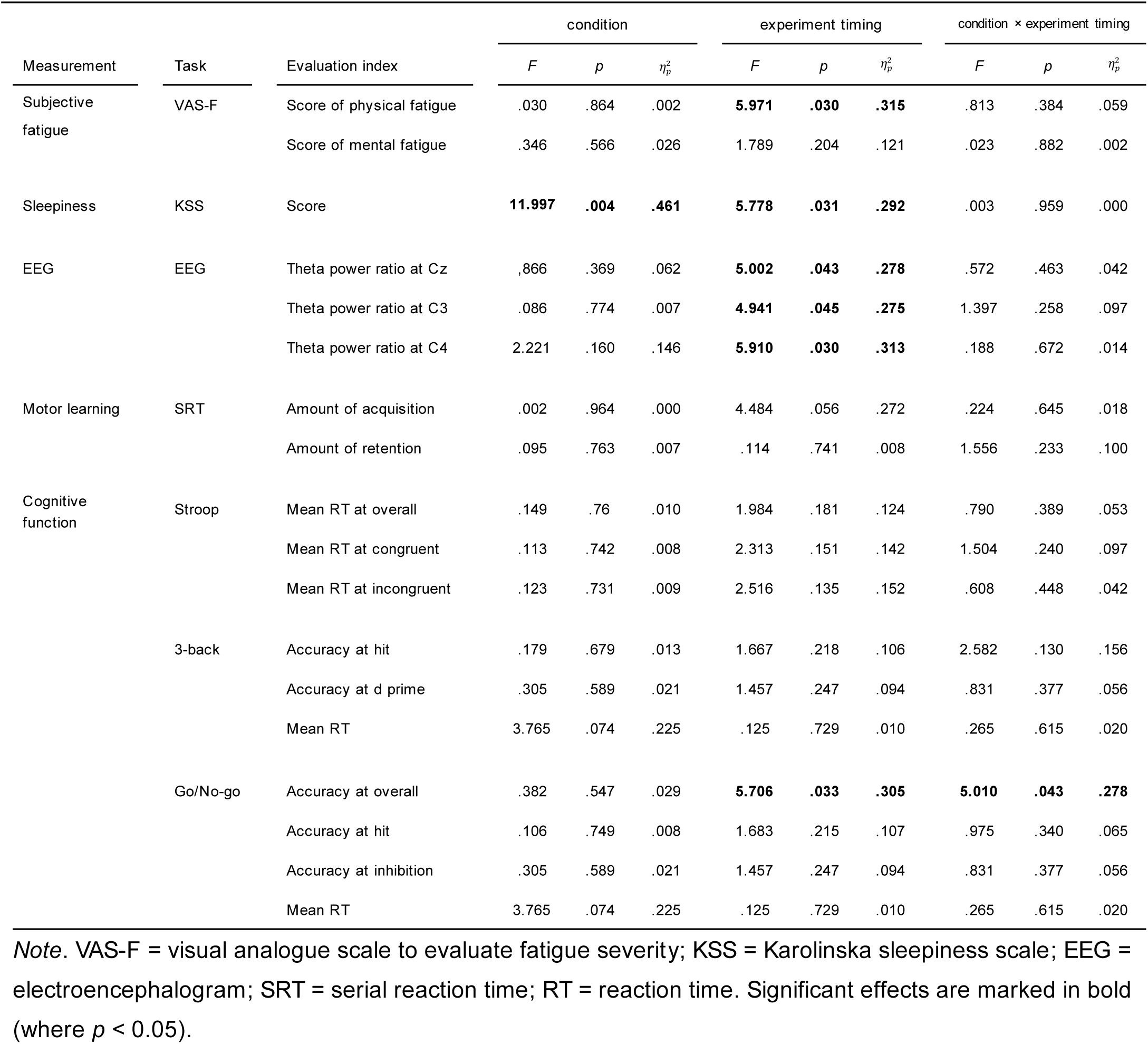
Results of two-way repeated-measures ANOVA for early chronotypes.

**Table 4:** Results of two-way repeated-measures ANOVA for late chronotypes.

| Measurement | Task | Evaluation index | condition |  |  | experiment timing |  |  | condition × experiment timing |  |  |
| --- | --- | --- | --- | --- | --- | --- | --- | --- | --- | --- | --- |
| | | | *F* | *p* | $\eta_p^2$ | *F* | *p* | $\eta_p^2$ | *F* | *p* | $\eta_p^2$ |
| Subjective fatigue | VAS-F | Score of physical fatigue | **5.133** | **.040** | **.268** | 2.263 | >.999 | .086 | 1.320 | .270 | .086 |
|  |  | Score of mental fatigue | 1.000 | .334 | .067 | 1.870 | .193 | .118 | .027 | .872 | .002 |
| Sleepiness | KSS | Score | 1.749 | .207 | .111 | .632 | .440 | .043 | .856 | .371 | .058 |
| EEG | EEG | Theta power ratio at Cz | .888 | .363 | .064 | .140 | .715 | .011 | .018 | .894 | .001 |
|  |  | Theta power ratio at C3 | .515 | .486 | .038 | .367 | .555 | .027 | .809 | .385 | .059 |
|  |  | Theta power ratio at C4 | .339 | .570 | .025 | .298 | .595 | .022 | .131 | .723 | .010 |
| Motor learning | SRT | Amount of acquisition | 1.749 | .207 | .111 | .632 | .440 | .043 | .856 | .371 | .058 |
|  |  | Amount of retention | .062 | .807 | .004 | .040 | .845 | .003 | .754 | .400 | .051 |
| Cognitive function | Stroop | Mean RT at overall | .047 | .831 | .003 | **32.461** | **<.001** | **.699** | .028 | .870 | .002 |
|  |  | Mean RT at congruent | .008 | .931 | .001 | **13.350** | **.003** | **.488** | .131 | .723 | .009 |
|  |  | Mean RT at incongruent | .047 | .831 | .003 | **32.461** | **<.001** | **.699** | .028 | .870 | .002 |
|  | 3-back | Accuracy at hit | .154 | .701 | .011 | 1.017 | .330 | .068 | 3.083 | .101 | .180 |
|  |  | Accuracy at d prime | .003 | .960 | .000 | 3.396 | .087 | .195 | .442 | .517 | .031 |
|  |  | Mean RT | .075 | .788 | .005 | .004 | .952 | .000 | **6.754** | **.021** | **.325** |
|  | Go/No-go | Accuracy at overall | 3.400 | .086 | .195 | 1.078 | .317 | .071 | .138 | .716 | .010 |
|  |  | Accuracy at hit | 2.266 | .154 | .139 | 1.345 | .265 | .088 | 1.386 | .259 | .090 |
|  |  | Accuracy at inhibition | 1.134 | .305 | .075 | .542 | .474 | .037 | **5.453** | **.035** | **.280** |
|  |  | Mean RT | 1.142 | .303 | .075 | .884 | .363 | .059 | .348 | .564 | .024 |

*Note.* VAS-F = visual analogue scale to evaluate fatigue severity; KSS = Karolinska sleepiness scale; EEG = electroencephalogram; SRT = serial reaction time; RT = reaction time. Significant effects are marked in bold (where $p < 0.05$ ).

#### KSS

To examine the influence of the time of day and schedule alignment on subjective sleepiness and alertness, participants’ subjective states were evaluated using the KSS. Lower scores represented reduced sleepiness. For ECs, the two-way rmANOVA results revealed significant main effects of both condition and experiment timing (*F* = 11.997, *p* = 0.004, 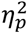 = 0.461; *F* = 5.778, *p* = 0.031, 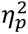 = 0.292, Figure 4, Table 3). For ECs, scores in the Evening were higher compared with those in the Morning, and scores in the Self-Time condition were higher compared with those in the Clock-Time condition (Figure 4, Table 1). For LCs, the two-way rmANOVA results revealed no significant main effects or interaction (Figure 4, Table 4).

This indicates that there was no significant difference between the Morning and Evening or between the Clock- and Self-Time conditions in LCs (Figure 4, Table 2). Taken together, the findings indicated that sleepiness was influenced by both the awake duration and the timing of sleep in ECs, whereas it showed no change in LCs.

### EEG

To assess physiological indicators of sleep pressure, the θ-band power ratio of EEG was evaluated with eyes open [23,41–44]. Higher power ratio values suggested an accumulation of sleep pressure [41–44]. For ECs, the two-way rmANOVA results revealed a significant effect of experiment timing (Cz, *F* = 5.002, *p* = 0.043, 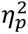 = 0.278; C3, *F* = 4.941, *p* = 0.045, 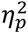 = 0.275; C4, *F* = 5.910, *p* = 0.030, 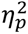 = 0.313, Figure 5, Table 3). The ratio at Cz, C3, and C4 was significantly lower in the Morning than in the Evening condition (Figure 5, Table 1). For LCs, the two-way rmANOVA results revealed no significant main effects or interaction (Figure 5, Table 4). This indicates that there was no significant difference between the Morning and Evening conditions or between the Clock- and Self-Time conditions in LCs (Figure 5, Table 2). These findings indicate that physiological sleep pressure in ECs showed clear time of day modulation, whereas LCs did not exhibit comparable changes across conditions.

**Figure 5:**
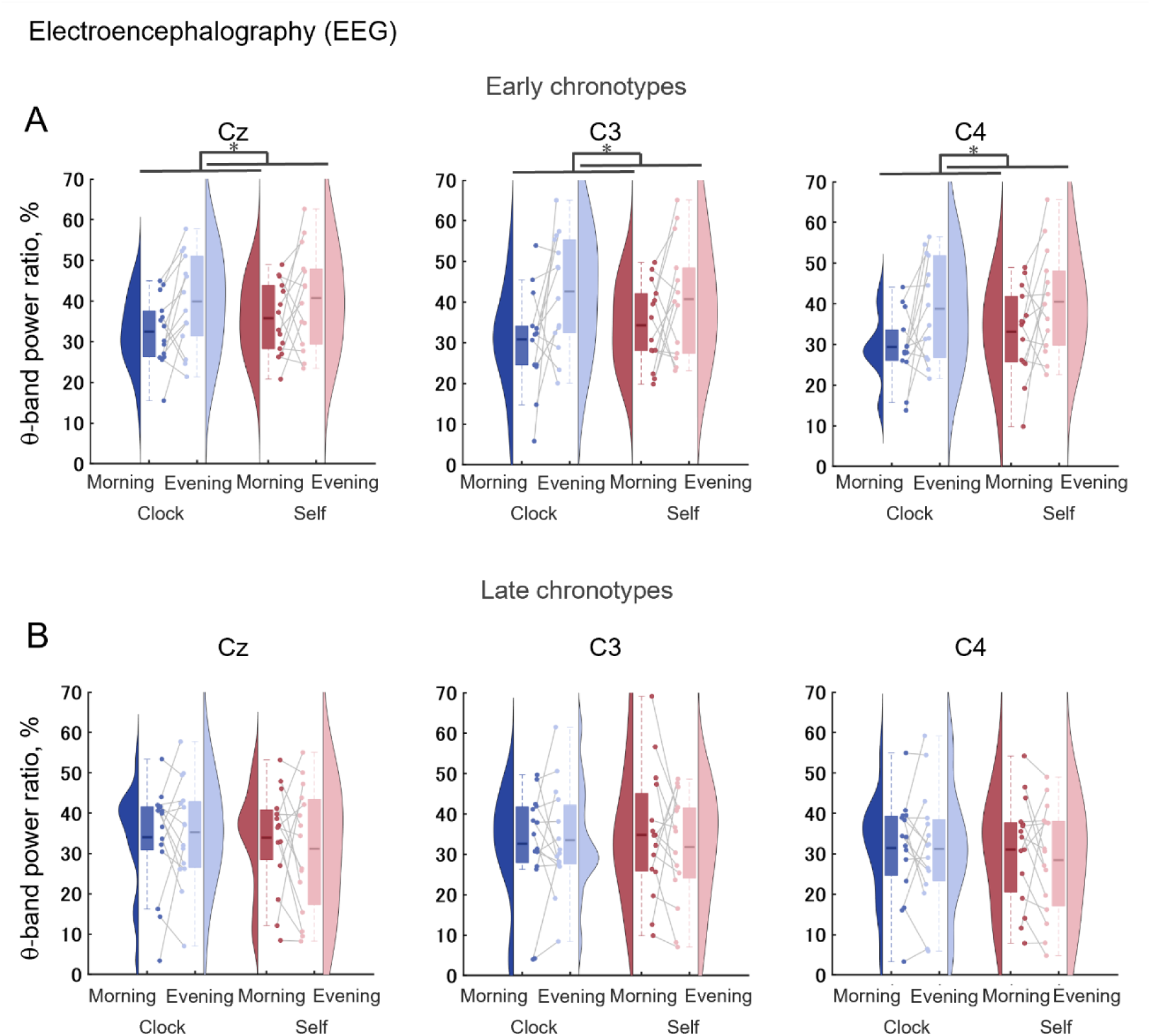
θ-band power ratio of electroencephalography (EEG) with eyes open in early and late chronotypes. θ-band power ratio of electroencephalography (EEG; sum of 4–7 Hz power relative to total 4–50 Hz power) with eyes open as indicators of sleep pressure was measured under Clock-Time conditions (Clock-Morning and Clock-Evening) and Self-Time conditions (Self-Morning and Self-Evening) in early chronotypes (ECs) and late chronotypes (LCs). The top panels (A) represent data for ECs and the bottom panels (B) represent data for LCs. **A,B** θ-band power ratio of EEG at Cz, C3 and C4. **A,B** were analyzed using a two-way repeated-measures analysis of variance to examine the effects of condition, experiment timing, and the interaction between condition and experiment timing on θ-band power ratio of EEG. Significant main effects of condition and experiment timing are indicated by asterisks (* for *p* < 0.05). Violin plots show the distribution of data. Boxplots show the mean and interquartile range. Dots and connecting lines indicate individual participant data.

### Behavioral measurements: a motor learning task and cognitive tasks

#### Motor learning task: SRTT

To evaluate motor learning efficiency, we assessed the amount of acquisition and retention using RTs in the SRTT. The difference in mean RT between blocks 5 and 6 was taken as an index of motor sequence learning acquisition, whereas the difference between blocks 6 and 7 was taken as an index of retention (see “Methods”). For ECs, about both the amount of acquisition and retention, the two-way rmANOVA results revealed no significant main effects or interactions (Figure 6, Table 3). This indicates that there was no significant difference between the Morning and Evening or between the Clock- and Self-Time conditions in ECs (Figure 6, Table 1). Similarly, for LCs, regarding both the amount of acquisition and retention, the two-way rmANOVA results revealed no significant main effects or interaction (Figure 6, Table 4). This indicates that there was no significant difference between Morning and Evening or between the Clock- and Self-Time conditions in LCs (Figure 6, Table 2). These findings suggest that motor learning acquisition and retention were preserved irrespective of chronotype and sleep-wake alignment.

**Figure 6:**
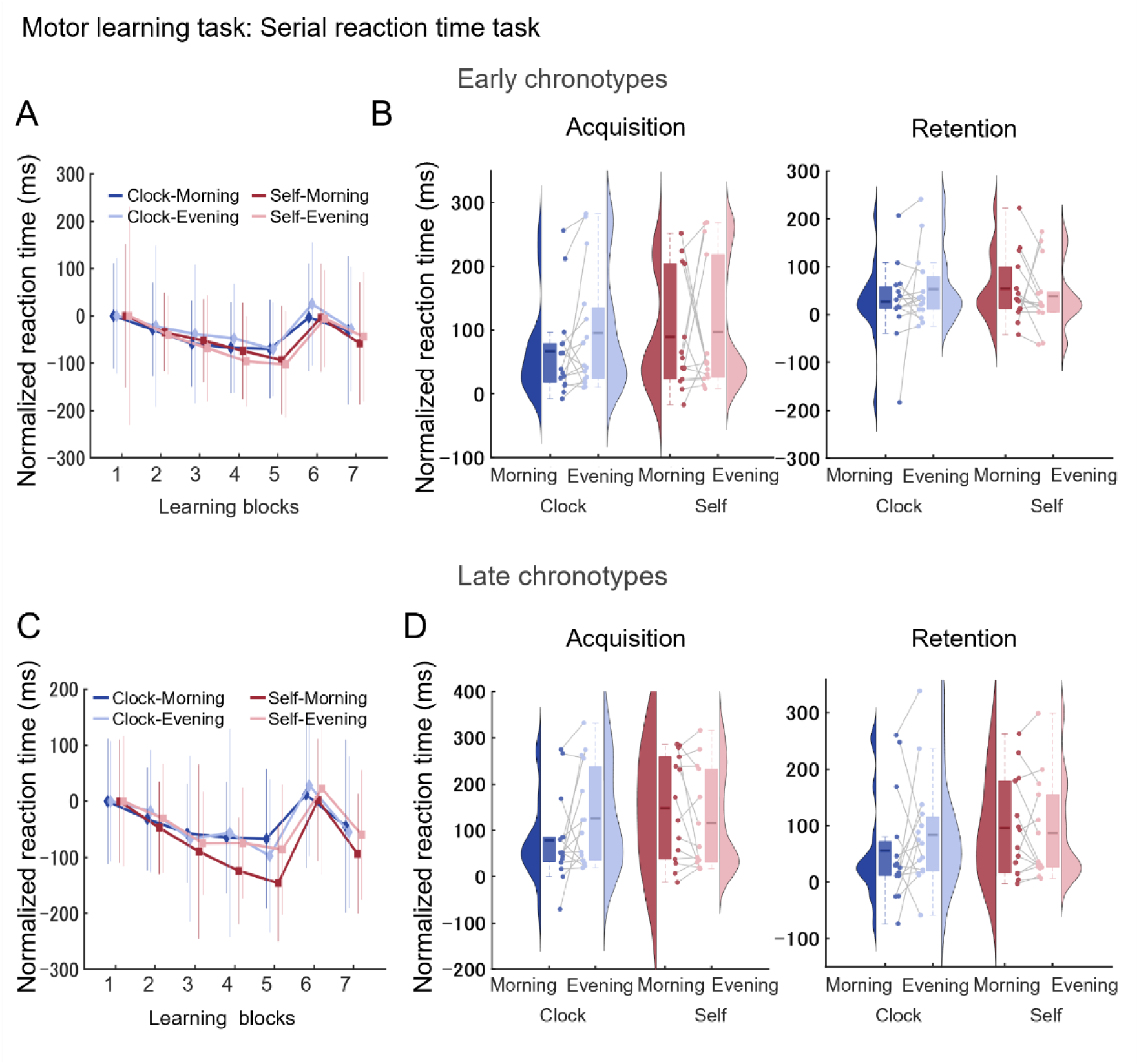
The amount of motor learning acquisition and retention for the serial reaction time task (SRTT) in early and late chronotypes. Reaction times (RTs) for the serial reaction time task (SRTT) were measured under Clock-Time conditions (Clock-Morning and Clock-Evening) and Self-Time conditions (Self-Morning and Self-Evening) in early chronotypes (ECs) and late chronotypes (LCs). The top panels (AB) represent data for ECs and the bottom panels (CD) represent data for LCs. **A, C** Overview of normalized mean RTs across blocks in each session. In block 1, the sequence followed a pseudorandom order. Mean RTs in block 1 served as the baseline of normalized RT. From blocks 2 to 5, the sequence of dots followed a fixed order, resulting in gradually faster RTs. In block 6, the sequence followed a pseudorandom order, leading to increased RTs. The differences in RTs between blocks 5 and 6 reflect the amount of acquisition, while the differences between blocks 6 and 7 indicate the amount of retention. **B, D** Shorter mean RTs indicate a lower amount of acquisition and retention. **B, D** were analyzed using a two-way repeated-measures analysis of variance to examine the effects of condition, experiment timing, and the interaction between condition and experiment timing on motor learning task. Violin plots show the distribution of data. Boxplots show the mean and interquartile range. Dots and connecting lines indicate individual participant data.

#### Cognitive tasks: Stroop color-word task

To evaluate selective attention, cognitive inhibition, and information processing speed, participants performed the Stroop color-word task. Performance was assessed by the mean RTs in all trials, congruent trials, and incongruent trials. Regarding mean RTs in all trials for ECs (overall, congruent, and incongruent), the two-way rmANOVA results revealed no significant main effects or interactions (Figure 7, Table 3). This indicates that there was no significant difference between Morning and Evening or between the Clock- and Self-Time conditions in ECs (Figure 7, Table 2). In contrast, regarding mean RTs in all trials in LCs (overall, congruent, and incongruent), the two-way rmANOVA results revealed a significant effect of experiment timing (*F* = 32.461, *p* < 0.001, 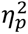 = 0.699; *F* = 13.350, *p* = 0.003, 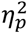 = 0.488; *F* = 32.461, *p* < 0.001, 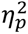 = 0.699, Figure 7, Table 4). The mean RTs were significantly smaller in the Evening than in the Morning in LCs (Figure 7, Table 2). These results indicated that the cognitive performance in the Stroop color-word task remained stable in ECs, whereas LCs showed faster responses in the later stage of awake duration with a chronotype-specific effect.

**Figure 7:**
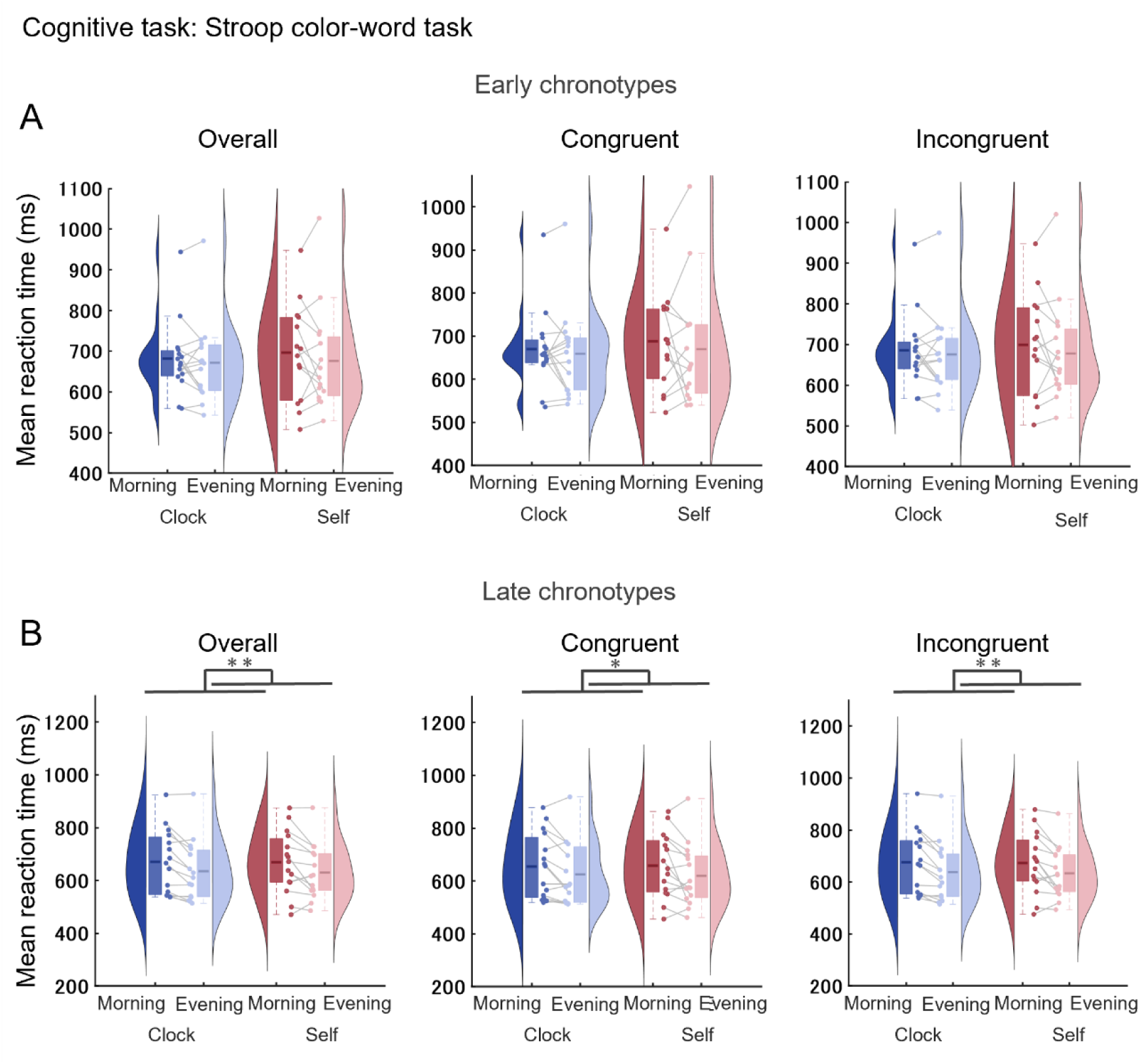
Reaction times for the Stroop color-word task in early and late chronotypes. Reaction times (RTs) for the Stroop color-word task were measured under Clock-Time conditions (Clock-Morning and Clock-Evening) and Self-Time conditions (Self-Morning and Self-Evening) in ECs and LCs. The top panels (A) represent data for ECs and the bottom panels (B) represent data for LCs. **A, B** Mean overall RTs, congruent trial RTs, and incongruent trial RTs for the Stroop color-word task. Smaller mean RTs indicate faster responses. **A, B** were analyzed using a two-way repeated-measures analysis of variance to examine the effects of condition and experiment timing and interaction between condition and experiment timing on the Stroop color-word task. Significant main effects of condition and experiment timing are indicated by asterisks (* for *p* < 0.05; ** for *p* < 0.01). Violin plots show the distribution of the data. Boxplots show the mean and interquartile range. Dots and connecting lines indicate individual participant data.

#### Cognitive tasks: 3-back letter task

To assess working memory performance, participants completed a 3 -back letter task. Performance was evaluated using accuracy measures (% hits and % d prime) and mean RTs. The sensitivity index *d* (or d prime) represents the difference between the hit rate and the false-alarm rate. For ECs, regarding the accuracy and mean RTs, the two-way rmANOVA results revealed no significant main effects or interaction (Figure 8, Table 3). This indicates that there was no significant difference between Morning and Evening or Clock- and Self-Time condition in ECs (Figure 8, Table 1). Similarly, for LCs, regarding accuracy, the two-way rmANOVA revealed no significant main effects or interaction (Figure 8, Table 4). Regarding mean reaction time, the two-way rmANOVA showed a significant interaction effect *F* = 6.754, *p* = 0.021, 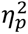 = 0.325, Figure 8, Table 4). However, neither the main effects of condition and experiment timing nor the simple main effects were significant (Figure 8, Table 4). This indicates that there was no significant difference between Morning and Evening or between the Clock- and Self-Time conditions in LCs (Figure 8, Table 2). These results indicated that working memory performance in the 3-back task was not influenced by the time of day or awake duration in both types.

**Figure 8:**
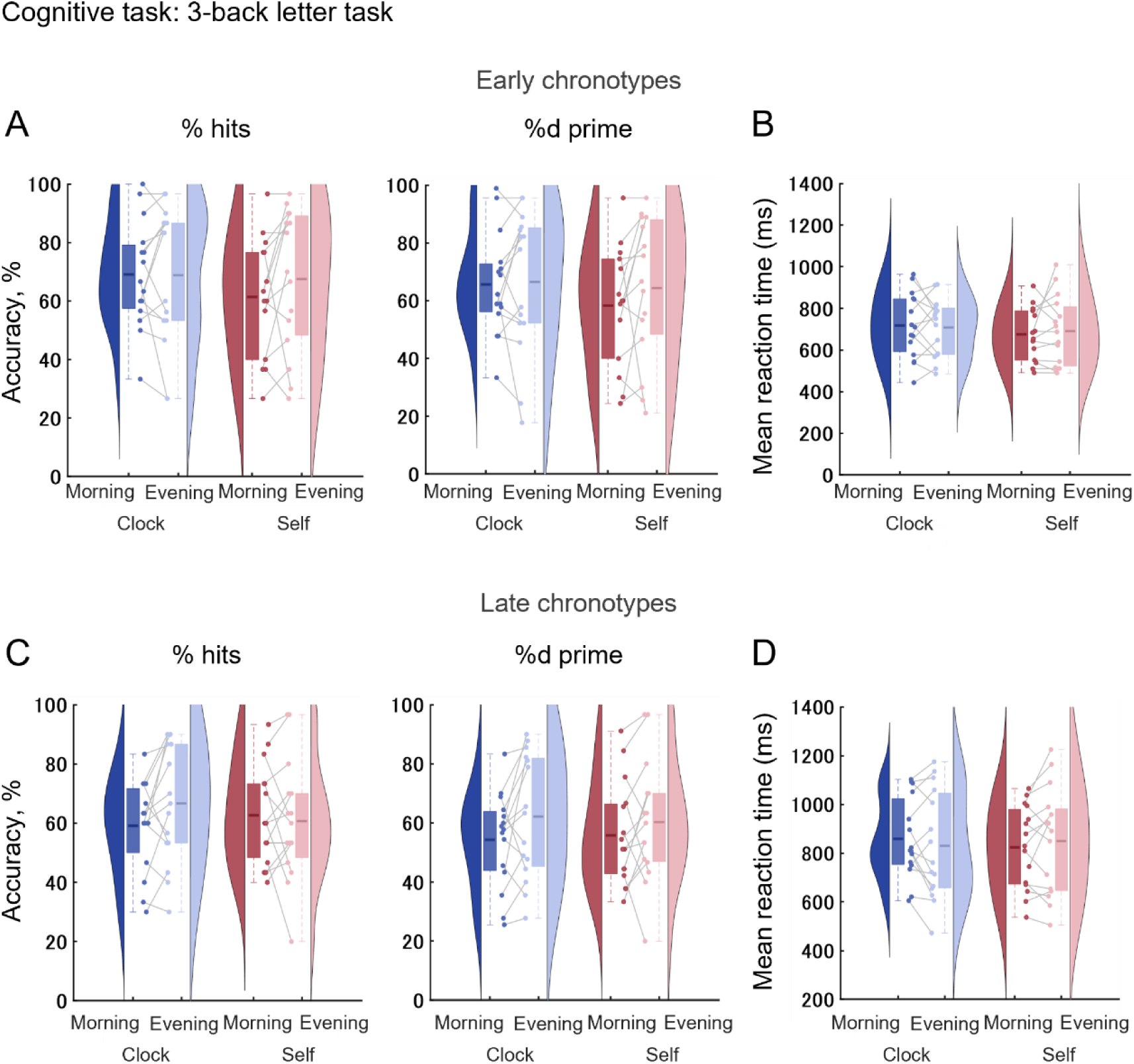
Accuracy and reaction times for the 3 -back letter task in early and late chronotypes. Accuracy and reaction times (RTs) for the 3-back letter task were measured under Clock-Time conditions (Clock-Morning and Clock-Evening) and Self-Time conditions (Self-Morning and Self-Evening) in ECs and LCs. The top panels (A, B) represent data for ECs and the bottom panels (C, D) represent data for LCs. **A, C** Accuracy of % hits and % d prime for the 3-back letter task. % hits represents the proportion of correct responses (hits) relative to the total number of trials. % d prime represents the proportion of hits minus the proportion of false-alarms. **B, D** Mean RTs for the 3-back letter task. Smaller mean RTs indicate faster responses. **A–D** were analyzed using a two-way repeated-measures analysis of variance to examine the effects of condition, experiment timing, and the interaction between condition and experiment timing on the 3-back letter task. Violin plots show the distribution of the data. Boxplots show the mean and interquartile range. Dots and connecting lines indicate individual participant data.

#### Cognitive tasks: Go/No-go task

To assess attentional functioning, including sustained and transient attention, as well as cognitive and impulse inhibition, the overall accuracy, hit trial accuracy, inhibition trial accuracy, and mean RTs in the Go/No-go task were evaluated. For ECs, regarding the accuracy of overall trials, the two-way rmANOVA showed a significant interaction effect (*F* = 5.010, *p* = 0.043, 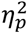 = 0.278, Figure 9, Table 1). Additionally, the results revealed no significant main effect of condition. However, a significant effect of experiment timing was observed (*F* = 5.706, *p* = 0.033, 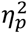 = 0.305, Figure 9, Table 1). The results showed a significant simple main effect on the Self-Time condition. Bonferroni-corrected post hoc comparisons indicated that the accuracy in the Self-Morning was significantly higher compared with that in the Self-Evening (Figure 9, Table 3). Regarding the accuracy of hit and inhibition trials and mean RTs, the two-way rmANOVA results revealed no significant main effects or interaction (Figure 9, Table 3). These results indicate that there was no significant difference between Morning and Evening or between the Clock- and Self-Time conditions in ECs (Figure 9, Table 1). In LCs, regarding the accuracy of inhibition trials, the two-way rmANOVA showed a significant interaction effect (*F* = 5.453, *p* = 0.035, 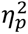 = 0.280, Figure 9, Table 4). However, there was no significant main effect of condition or experiment timing (Figure 9, Table 4). The results revealed a significant simple main effect on the Clock-Time condition. Bonferroni-corrected post hoc comparisons indicated that the score in the Clock-Morning condition was significantly lower compared with that in the Clock-Evening condition (Figure 9, Table 2). Regarding the overall accuracy, the accuracy of hit trials, and mean RTs, the two-way rmANOVA results revealed no significant main effects or interaction (Figure 9, Table 4). This result indicated that there was no significant difference between Morning and Evening or between the Clock- and Self-Time conditions in LCs (Figure 9, Table 2). Taken together, the findings indicated that Go/No-go task performance was influenced by awake duration. ECs showed better performance during the earlier stage of wakefulness, whereas LCs performed better during the later stage.

**Figure 9:**
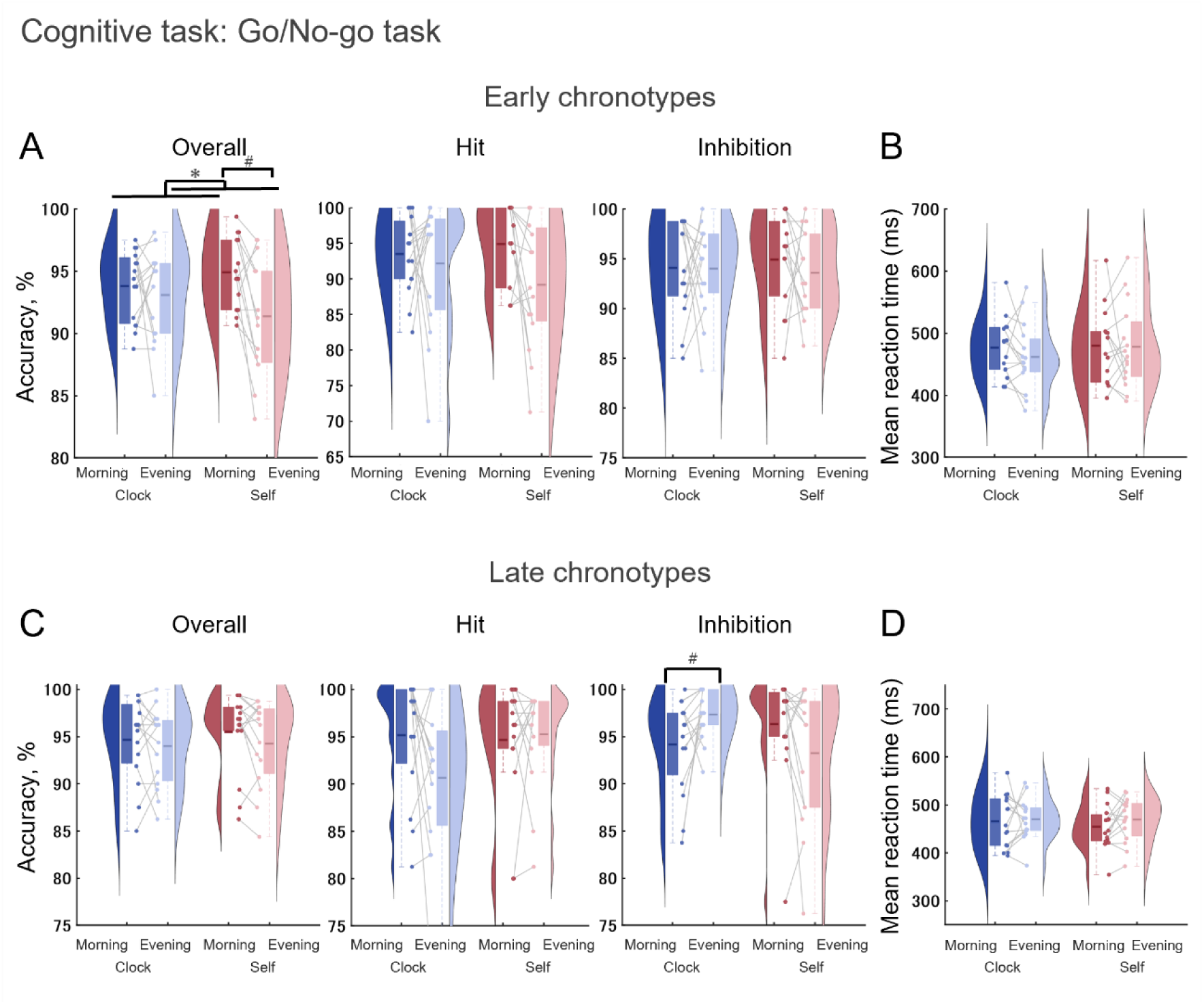
Accuracy and reaction times for the Go/No-go task in early and late chronotypes. Accuracy and rection times (RTs) for the Go/No-go task were measured under Clock-Time conditions (Clock-Morning and Clock-Evening) and Self-Time conditions (Self-Morning and Self-Evening) in ECs and LCs. The top panels (A, B) represent data for ECs, and the bottom panels (C, D) represent data for LCs. **A, C** Overall accuracy, hit trial accuracy, and inhibition trial accuracy for the Go/No-go task. The overall accuracy is the proportion of correct responses (hits) relative to the total number of trials. Hit trial accuracy is the proportion of correct responses for “Go” trials relative to the total number of “Go” trials. Inhibition trial accuracy is the proportion of correct responses for “No-go” trials relative to the total number of “No-go” trials. **B, D** Mean RTs in hit trials for the Go/No-go task. Smaller mean RTs indicated faster responses. **A–D** were analyzed using a two-way repeated-measures analysis of variance to examine the effects of condition and experiment timing and interaction between condition and experiment timing in the Go/No-go task. Significant main effects of condition and experiment timing are indicated by asterisks (* for *p* < 0.05; ** for *p* < 0.01). Bonferroni-corrected post hoc tests revealing significant differences are indicated by hashes (# for *p* < 0.05). Violin plots show the distribution of data. Boxplots display the mean and interquartile range. Dots and connecting lines indicate individual participant data.

## Discussion

In the present study, we aimed to clarify how aligning or misaligning sleep-wake schedules with individual chronotypes influences daily variations in subjective, behavioral, and physiological parameters. Through comparisons between two conditions (sleep aligned with social rhythms [Clock-Time condition] and sleep aligned with individual preferences [Self-Time condition]), we found that sleep timing influenced physiological and psychological parameters, whereas behavioral parameters were largely independent of it. Indeed, regarding subjective state, ECs showed increases in fatigue and sleepiness with awake duration, reflecting typical accumulation of sleep pressure, whereas LCs were more affected by sleep timing and exhibited atypical accumulation patterns. Motor learning tended to improve later in the day in both chronotypes, regardless of sleep timing. In contrast, cognitive performance showed different peaks in each chronotype determined by awake duration -the earlier stage for ECs and the later stage for LCs-regardless of sleep timing. These findings highlight the potential to develop strategies for improving human behavior and physiology by considering both sleep timing and chronotype, and demonstrate that performance optimization requires understanding of the complex interplay between circadian rhythms and awake duration.

### Sleep timing as a key determinant of fatigue and sleepiness

These findings revealed that both ECs’ and LCs’ subjective state was influenced by condition, whereas ECs’ sleepiness was affected by condition and awake duration. Inadequate sleep duration is widely recognized as a significant public health concern [46] and substantial research attention has focused on the effects of the length of sleep. Despite identical sleep durations across conditions, the current results revealed that fatigue and sleepiness patterns were strongly influenced by sleep timing. Thus, these findings emphasize the importance of not only sleep duration but also sleep timing in managing physiological and psychological states.

### Chronotype-specific dynamics of sleep pressure with EEG

θ-band power ratio, an EEG marker of sleep pressure, typically increases with an increasing duration of time awake [43]. However, in the current study, we observed marked chronotype-dependent differences in this accumulation pattern. In ECs, the θ-band power ratio in the later portion of wakefulness was higher compared with that in the earlier portion of wakefulness, whereas LCs showed no such increase. This atypical pattern in LCs is likely to contribute to their difficulty in sleep initiation compared with ECs’ relative ease, and potentially explains their delayed sleep-wake cycle. While current interventions for aligning LCs to social schedules are primarily focused on light therapy and pharmacological approaches [11,12], our findings suggest that addressing the rate of sleep pressure accumulation may provide a novel and potentially effective therapeutic pathway. Specifically, neurofeedback training protocols designed to enhance the progressive increase of θ-band power throughout the waking period may be helpful for normalizing sleep pressure dynamics in LCs. Such interventions could be useful for training LCs to develop the typical pattern of sleep pressure buildup that occurs with sustained wakefulness, potentially facilitating earlier sleep initiation and improving circadian alignment with social demands. When integrated with traditional circadian-focused strategies, these neurophysiology-based approaches could provide a comprehensive framework for mitigating circadian misalignment and supporting LCs in adapting to conventional social schedules.

### Motor learning tends to improve in the afternoon regardless of sleep timing across chronotypes

Motor learning performance tends to be consistently enhanced later in the day, such as in the afternoon and evening, regardless of chronotype in the current study. For ECs, the amount of motor learning acquisition tended to be greater at the Clock-Evening (07:00 PM) and Self-Evening (approximately 08:30 PM) time points, whereas for LCs, it tended to be greater at the Clock-Evening (07:00 PM) and Self-Morning (01:00 PM) time points. These patterns may be better understood by considering the sleep timing at each condition between chronotypes. For ECs, the timing of the Self-Time condition closely resembled that of the Clock-Time condition, because ECs tended to follow conventional nighttime sleep schedules (10:30 PM–06:30 AM). For ECs, Self-Morning occurs in the actual morning hours, and Self-Evening aligns with their natural evening, at approximately 08:30 PM. In contrast, LCs, who typically go to bed and wake up much later than ECs, showed a marked delay in the experiment timing of the Self-Time condition. For LCs, Self-Morning (measured 1.5 hours after waking) occurs in the afternoon, while Self-Evening (measured 12.5 hours after waking) corresponds to nighttime. This indicates that, despite differences in sleep timing and awake duration between chronotypes, motor learning acquisition tended to improve during the afternoon and evening hours in both chronotypes. These findings suggest that the time of day exerts an influence on motor learning regardless of individual sleep schedules and awake duration.

This enhancement of motor learning in the afternoon and evening aligns with neurophysiological research on the effects of the time of day on cortical excitability in the primary motor cortex. Transcranial magnetic stimulation studies have reported that cortical inhibition is higher in the morning [47] and cortical excitability increases in the evening [48]. Enhanced cortical excitability is associated with improved motor learning [23]. Considering physiological data from these previous studies and the motor learning results from the present study together suggests that motor learning is enhanced during evening hours when cortical excitability is elevated. Previous studies conducted under conventional nighttime sleep schedules have been unable to determine whether cortical excitability is enhanced specifically in the evening or during the latter part of the awake duration. However, the results of the current study, with varying sleep schedules, suggest that evening time itself is crucial for motor learning enhancement rather than the awake duration.

The current findings may have important practical implications, particularly in the fields of sports training and rehabilitation. Regarding sports training, the results suggest that activities requiring precise motor skill acquisition, such as mastering dance choreography or developing technical sport-specific skills, are likely to achieve better results when scheduled during the afternoon or evening. Similarly, regarding rehabilitation, our findings suggest that scheduling therapeutic interventions in the afternoon or evening might enhance recovery outcomes by leveraging periods of heightened motor learning capacity. Such timing optimization could greatly benefit activities in which motor learning is critical for performance enhancement.

### Cognitive performance peaks depend noton the absolutetime of day but on the awake duration

Although motor learning tends to be affected by time of the day, it should be noted that cognitive performance appeared to be affected by awake duration, irrespective of sleep timing. Specifically, ECs exhibited enhanced cognitive performance during the earlier stages of awake duration. In contrast LCs showed improvements during the latter stages of awake duration in both the Clock- and Self-Time conditions. Most previous studies of this topic have reported that ECs perform better in the morning and LCs perform better in the evening, attributing these results to circadian influences [21–27]. However, these previous studies were conducted under conventional nighttime sleep schedules, making it difficult to determine whether the observed effects reflected intrinsic influences of the time of day or the awake duration. In contrast, in the present study, performance was examined under both a conventional nighttime sleep schedule (the Clock-Time condition) and a self-selected sleep schedule aligned with participants’ chronotype preferences (the Self-Time condition). Using the present research protocol, we were able to isolate the influence of waking duration from the clock time.

These results revealed that the observed differences in performance were not solely attributable to fixed clock times during the day, but were instead modulated by the awake duration from each individual’s wake-up time. This finding challenges the conventional assumption that cognitive and motor performance are determined by specific times of day (e.g., morning or evening), and highlights the need to consider waking duration as a key factor in understanding performance variations. This insight has important implications for chronotype-specific considerations in educational, occupational, and clinical contexts. By shifting the focus from conventional time of day models to awake duration and individual sleep-wake patterns, it may be possible to promote better performance across diverse chronotypes.

### The importance of naturalistic measurements in chronotype research

The present findings regarding the optimal performance timing for ECs under the Clock-Time condition diverged from the findings of traditional chronotype studies, which have reported that ECs exhibit better performance during the morning hours in both motor learning and cognitive tasks [21–27]. In contrast, the present findings demonstrated that ECs’ motor learning performance was enhanced during the afternoon, and some cognitive performance patterns varied across individuals. These discrepancies may arise from fundamental differences in experimental settings. Traditional chronotype studies have typically been conducted in highly controlled laboratory environments. While such settings ensure rigorous control of experimental variables, they limit ecological validity. In contrast, the current study adopted a naturalistic approach by assessing participants in their daily life envir onments, enabling us to capture authentic physiological responses in real-world contexts.

This methodological shift yielded two significant insights into chronobiology and research methodology. First, our findings suggest a difference in sensitivity to timing effects between chronotypes. LCs consistently exhibited strong performance in the evening, even in less controlled environments. However, ECs displayed individual variability in certain cognitive tasks, and their responses to the time of day reflected greater flexibility in optimal performance timing than previously assumed. This variability indicates that EC performance patterns are more context-dependent and individually variable compared with LC performance patterns, with LCs consistently preferring evenings. These findings introduce a new perspective on chronotype differences, suggesting that beyond simple timing preferences, chronotypes may differ in their underlying sensitivity to temporal factors. Second, our findings highlight the importance of investigating brain function and behavior in naturalistic environments, in accord with the growing recognition of the need for “making experiments more natural in order to decode the brain” [49]. Using this wearable EEG system, we were able to bridge the gap between ecological validity and precise physiological measurement, addressing the classic “lab dilemma” in psychological research, which reflects the trade-off between experimental control and the ability to capture authentic behavior [50–52]. Real-world settings allow researchers to capture individuals’ performance under authentic conditions. This is particularly crucial in chronobiology research, in which environmental factors can influence performance.

### Study limitations

Although the current study provides valuable insights into the relationships among sleep timing, circadian rhythms, subjective state, sleep pressure, and behavior, several limitations should be acknowledged.

First, although the current study examined both motor learning and cognitive performance, it did not disentangle the underlying perceptual and neural mechanisms that might influence both processes. A deeper understanding of how these shared perceptual mechanisms evolve over time will be necessary to elucidate the precise relationships between motor learning and cognitive function. Future studies should directly capture neural activity using EEG and transcranial magnetic stimulation during task performance to elucidate the perceptual mechanisms involved. This approach could help clarify the contributions of sensory, cognitive, and motor processes to these domains and refine our understanding of their interactions.

Second, the timing of measurements represents a key limitation of the current study. Following previous research [23], measurements were conducted in the morning (at 08:00 AM, 1.5 hours post-awakening) and in the evening (at 07:00 PM, 12.5 hours post-awakening). Given the burden on participants, it was not feasible to collect data at multiple additional time points within a day. This limitation is particularly significant given the present finding that LCs exhibited atypical patterns of sleep pressure accumulation. Previous studies have highlighted the dynamic nature of alertness and its dependency on factors such as prior sleep, physical activity, and glucose levels throughout the day [53]. Future research should include measurements taken immediately before bedtime to provide a more comprehensive understanding of how parameters such as subjective states, sleep pressure, and performance measures fluctuate across the day and influence sleep initiation. Collecting these data in LCs would be helpful for clarifying the relationship between their unique sleep pressure dynamics and other physiological and behavioral parameters near sleep onset. This may provide critical insights into the mechanisms that govern sleep-wake regulation in LCs.

## Conclusion

The findings of the current study underscore the need to consider both sleep timing and chronotypes when examining psychological states, motor learning, cognitive performance, and sleep pressure. ECs exhibited steady fatigue accumulation, whereas LCs were more affected by misalignment with preferred sleep timing, with EEG data suggesting impaired sleep pressure dynamics. The results revealed that motor learning improved later in the day across chronotypes, whereas cognitive performance depended on awake duration. These findings emphasize that behavioral, physiological, and psychological dynamics are shaped by the interaction between chronotype and sleep timing, highlighting the importance of further research integrating circadian and sleep-wake schedule perspectives to advance our understanding of human health and functioning.

## Acknowledgments

This work was supported by Taikichiro Mori Memorial Research Grants to T.H, Keio Gijuku Academic Development Funds to J. Ushiyama, and a donation from Living Platform, Ltd., Japan to J. Ushiyama. We thank Ms. Tomomi Hamaoka for their secretarial assistance and all members of our laboratory for their insightful comments on the work. We thank Benjamin Knight, MSc., from Edanz (https://jp.edanz.com/ac) for editing a draft of this manuscript.

